# Assessing different components of biodiversity across a river network using eDNA

**DOI:** 10.1101/546549

**Authors:** Elvira Mächler, Chelsea J. Little, Remo Wüthrich, Roman Alther, Emanuel A. Fronhofer, Isabelle Gounand, Eric Harvey, Samuel Hürlemann, Jean-Claude Walser, Florian Altermatt

## Abstract

1. Assessing individual components of biodiversity, such as local or regional taxon richness, and differences in community composition is a long-standing challenge in ecology. It is especially relevant in spatially structured and diverse ecosystems. Environmental DNA (eDNA) has been suggested as a novel technique to accurately measure biodiversity. However, we do not yet fully understand the comparability of eDNA-based assessments to previously used approaches.
2. We sampled may-, stone-, and caddisfly genera with contemporary eDNA and kicknet methods at 61 sites distributed over a large river network, allowing a comparison of various diversity measures from the catchment to site levels and providing insights into how these measures relate to network properties. We extended our survey data with historical records of total diversity at the catchment level.
3. At the catchment scale, eDNA and kicknet detected similar proportions of the overall and cumulative historically documented species richness (gamma diversity), namely 42% and 46%, respectively. We further found a good overlap (62%) between the two contemporary methods at the regional scale.
4. At the local scale, we found highly congruent values of local taxon richness (alpha diversity) between eDNA and kicknet. Richness of eDNA was positively related with discharge, a descriptor of network position, while kicknet was not.
5. Beta diversity between sites was similar for the two contemporary methods. Contrary to our expectation, however, beta diversity was driven by species replacement and not by nestedness.
6. Although optimization of eDNA approaches is still needed, our results indicate that this novel technique can capture extensive aspects of gamma diversity, proving its potential utility as a new tool for large sampling campaigns across hitherto understudied complete river catchments, requiring less time and becoming more cost-efficient than classical approaches. Overall, the richness estimated with the two contemporary methods is similar at both local and regional scale but community composition is differently assessed with the two methods at individual sites and becomes more similar with higher discharge.

## Introduction

Quantifying biodiversity accurately is a long-standing challenge of primary importance in ecology (Dornelas et al., 2013, Gotelli and Colwell, 2001, Whittaker, 1972). On the one hand, there is a fundamental interest to understand the distribution of diversity in time and space and the mechanistic drivers from local to regional scales (e.g., Gaston, 2000, Gotelli and Colwell, 2011, Koleff et al., 2003). On the other hand, the state of ecosystems is inherently linked to biodiversity (Pennekamp et al., 2018) and the current loss of biodiversity has potentially large negative consequences on the functions and services of ecosystems (Chapin et al., 2000, Isbell et al., 2017). This is especially relevant for freshwater habitats because they provide crucial ecosystem services such as drinking water, food security, or recreational value to humanity (Cardinale et al., 2012, Dudgeon et al., 2006, Postel and Carpenter, 1997).

In river systems, may-, stone-, and caddisflies (Ephemeroptera, Plecoptera, and Trichoptera; thereafter abbreviated as EPT) are often used as indicators due to their sensitivity to environmental change, their different preferences for ecological niches, and their relatively well-known taxonomy (Schmidt-Kloiber and Hering, 2015). Presence or absence of certain EPT taxa or their overall richness is highly informative and can be tightly linked to habitat quality. Importantly, they describe not only the current state of a water body but also integrate its changes over time. Thus, EPT are at the heart of many freshwater quality assessments around the world and are included in many regulatory frameworks, such as the Water Framework Directive (Directive 2000/60/EC, but see also Borja et al., 2009), or the Canadian Aquatic Biomonitoring Network (CABIN, Reynoldson et al., 2003).

Classically, EPT are collected with a standardized kicknet method (Barbour et al., 1999). Taxa are then identified under a dissecting microscope, which is time-consuming and therefore costly. Taxon richness is the most fundamental approach to estimate biodiversity and is still widely used. Even though the number of taxa is a convincingly intuitive proxy of biodiversity and the basis of many fundamental concepts in ecology, it is a difficult variable to measure accurately (Gotelli and Colwell, 2011, Purvis and Hector, 2000). Diversity can be further divided into different components, such as local richness, regional richness and between-site dissimilarity (also known as alpha, gamma, and beta diversity). The emerging technique of environmental DNA (eDNA) metabarcoding is expected to become a complementary or even replacement method (for example see Baird and Hajibabaei, 2012, Deiner et al., 2017, Lawson Handley, 2015). About a decade ago, the first study was published on the detection of species through DNA in an environmental sample (Ficetola et al., 2008). With the implementation of high throughput sequencing, which allowed not only detection of single species but whole communities, eDNA metabarcoding has been proposed to revolutionize biodiversity assessments (Bohmann et al., 2014, Lawson Handley, 2015, Shokralla et al., 2012). As with every novel technique, eDNA metabarcoding creates new opportunities but also challenges, especially in terms of recognizing what information it provides and how it compares to previously implemented and established methodologies.

Comparisons between eDNA and traditional methods have hitherto mostly focused on either local or regional richness comparisons. In river systems, previous studies have generally detected higher taxon richness with eDNA than kicknet approaches (Civade et al., 2016, Olds et al., 2016, Valentini et al., 2016). Those results were likely due to eDNA at one location integrating taxon information from upstream reaches via downstream transportation of eDNA (Deiner and Altermatt, 2014, Deiner et al., 2016, Pont et al., 2018). This suggests that standard techniques represent a more accurate local estimate while eDNA integrates information across space. In that context, any comparison of the two techniques will be influenced by the scale of the study. However, we still do not know if findings are directly comparable, complementary, or different across different spatial scales and across different components of biodiversity, due to the specific properties of kicknet and eDNA sampling. A detailed understanding is needed to make decisions on how to sample biodiversity in complex landscapes and to compare both local and regional measures. This is particularly relevant in river landscapes, where the typical underlying dendritic network structure is known to affect biodiversity (Altermatt, 2013, Altermatt et al., 2013, Carrara et al., 2012, Harvey et al., 2018, Tonkin et al., 2018).

In our study, we compared different measures of biodiversity of EPT sampled classically (i.e., by kicknet) or by eDNA, using a spatially structured approach that representatively covered a river network in a 740 km^2^ catchment. We analyzed how these two different approaches capture the facets of biodiversity at the level of alpha (local site), beta (between sites), and gamma diversity (catchment level). Gamma diversity information was supplemented with all historically available data. Given that taxon richness is among the most-commonly studied biodiversity variables, we discuss the design of biodiversity monitoring with eDNA in dendritic river networks.

## Material & Methods

We studied a river network in a 740 km^2^ catchment containing 61 sampling sites in the upper river Thur in north-eastern Switzerland (Fig.1, Table S1). The catchment comprises three main river-stems: Thur, Glatt, and Necker, the latter two draining into the Thur. In a sampling campaign conducted from June 11 to June 22 2016, we collected eDNA and benthic invertebrate kicknet samples, with the two sampling methods performed at each site within a two-day window. To characterize the position of sites in the network, we extracted stream order, catchment area and the mean annual discharge data for each site from existing databases (BAFU, 2013, BAFU, 2014, Faundler et al., 2013).

**Fig. 1:**
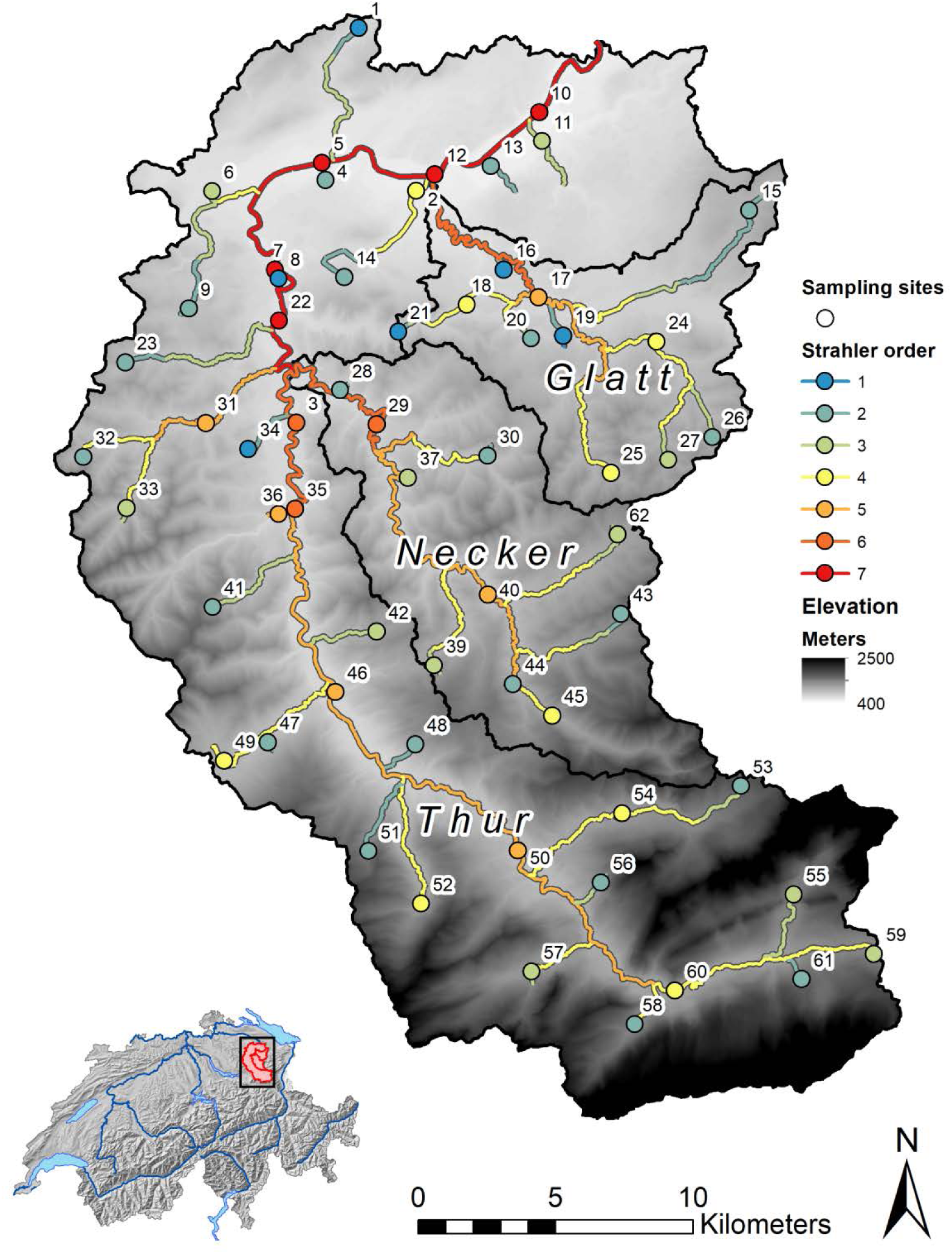
Field sites in the river Thur catchment in North-Western Switzerland (insert at bottom right). Colors are coding for stream order (first to seventh order streams: blue, green, yellow, orange, orange-red, and red respectively). Black lines indicate the three major subcatchments: Thur, Necker and Glatt. Data source: swisstopo: VECTOR200 (2017), DEM25 (2003), SWISSTLM3D (2018); BAFU: EZG (2012); Bundesamt für Landestopographie (Art.30 Geo IV): 5704 000 000, reproduced by permission of swisstopo / JA100119.

### Historical data

We obtained long-term biodiversity data on gamma diversity in our catchment from the Centre Suisse de la Cartographie de la Faune (CSCF). These data include all EPT species ever recorded in the whole catchment over the time-period 1981–2016. The data are of various sampling origins, but of high quality, thus giving a highly robust and reliable cumulative estimate of EPT genus richness (and the respective EPT genus identity) at the whole catchment scale. The cumulative data consist of 3,467 individual records based on observations of the species in an area of the catchment, which we then converted into genus richness.

### Contemporary kicknet data

We collected benthic macroinvertebrates based on three-minute kicknet sampling applied to three microhabitats present at a given site (Barbour et al., 1999). Leaves and debris were removed from the sample and the remaining material was pooled and stored in 96% molecular grade ethanol. In the lab, all EPT individuals were identified with a microscope to species level. A few taxa, only present as early instar larva or containing cryptic species, were grouped in pre-defined complexes, subsequently treated at the genus. We could not assess EPT taxa at one site due to the loss of a sample.

### eDNA filtration in the field

At each site, we sampled three times 250 mL of river water, each on a separate filter (for detailed description see Mächler et al., 2018). We collected eDNA samples about 5–10 meters upstream of the kicknet sampling to minimize cross-contamination. Samples were stored in a Styrofoam box equipped with cooling elements until we came back from the field (no longer than 9 h). Thereafter, they were stored at −20 °C until further processing. On each field day, we performed a replicated filter control (FC) that was filtered in the field before any sampling site was visited in order to check if the reusable material was clean. In total, we generated 11 filter controls, each consisting of three replicates. Operational taxonomic units found in at least two replicates of a filter control from the same date were removed from all samples for the further analysis. Further information on filtration, eDNA facilities, and material preparation can be found in the supplementary file.

### Extraction and library preparation

Detailed information about the extraction and library preparation can be found in the supplementary file. In short, we used the DNeasy Blood and Tissue kit (Qiagen GmbH, Hilden, Germany) to extract the DNA from eDNA samples, also including extraction controls (EC). We used an Illumina MiSeq dual-barcoded two-step PCR amplicon sequencing protocol. The first PCR was performed with modified primers that contained an adaptor-specific tail, a heterogeneity spacer, and the amplicon target site (see Table S2). On each of the PCR plates, we implemented one negative (NC) and one positive PCR control (PC). The negative control consisted of 5 μL sigma water and the positive control consisted of 4 μL sample and 1 μL (0.01 ng/μL) artificial dummy DNA (a randomly generated double-stranded DNA sequence that was 313 bp in length and matched primer region, see supplementary information). We then pooled the three PCR replicates per sample and cleaned it with SPRI beads. For the second PCR, we used the Nextera XT Index kit v2 (Illumina California, USA) to index each sample and cleaned afterwards the index reaction with SPRI beads. We quantified each sample with the Spark 10M Multimode Microplate Reader (Tecan Group Ltd., Männedorf, Switzerland) and pooled them in equimolar parts into a final pool that we cleaned with SPRI beads. All controls (FC, EC, PC, NC) were run alongside the samples and were pooled according to their concentrations. Controls that were too low to quantify were pooled into the second lowest concentrated pool with 10 µL, equal to the volume of the lowest sample in the respective pool. The libraries were added at 16 pM concentration and PhiX control was added at a 10% concentration. A paired-end (2×300 nt) sequencing was performed on an Illumina MiSeq (MiSeq Reagent kit v3, 300 cycles) following the manufacture’s run protocols (Illumina, California, USA). To increase the sequencing depth, a second run of the same pooled libraries was conducted.

### Bioinformatic data processing

After the two successful Illumina MiSeq runs, the data was demultiplexed and the quality of the reads was checked with FastQC (Andrews, 2010). Raw reads were end-trimmed (usearch v10.0.240, R1:30nt, R2:50nt) and merged with an overlap of min 15 bp max 300 bp (Flash, v1.2.11). Next, the primer sites were removed (full length, no mismatch allowed (cutadapt v1.12)) and thereafter, the data was quality filtered (prinseq-lite v0.20.4) using the following parameters: size range (100–500), GC range (30– 70), mean quality (20), and low complexity filter dust (30). In a next step, UNOISE3 (usearch v10.0.240) was used to determine amplicon sequence variants (zero-radius OTUs, thereafter called ZOTUs). UNOISE3 has a build-in error-correction to reduce the influence of sequencing errors (Edgar, 2016). An additional clustering at 99% sequence identity was performed to reduce sequence diversity and to account for possible amplification errors in the first PCR. The resulting ZOTUs (zero-radius OTUs, thereafter called ZOTUs) were checked for stop codons using the invertebrate mitochondrial code, to ensure an intact open reading frame. This resulted in 27 M reads corresponding to 11,313 ZOTUS (Table S3). As a final step, the ZOTUs were assigned to taxa (blast 2.3.0 and usearch v10.0.240, tax filter = 0.9).

### Statistics

Our analysis was based on the following strategy: First, we confirmed that the two Illumina runs could be combined and second we cleaned the sequencing data and selected only ZOTUs that were assigned to an EPT order (see detailed information in the supplementary file). For individual sites, ZOTUS were only counted if they were present in at least two of the three independent replicates (Fig. S1), which is a highly stringent assumption and conservative with respect to detection of taxa. Thereafter, we were able to analyze various diversity measures to identify similarities and differences between eDNA and kicknet sampling approaches: (i) gamma, (ii) alpha, (iii) beta diversity, and (iv) between method community dissimilarity. All statistical analyses were performed using R (R Core Team, version 3.4.4).

#### Gamma diversity

We used the R package ‘venneuler’ (Wilkinson and Urbanek, 2011) to draw Venn diagrams for gamma richness of historic, eDNA and kicknet data. We used the R package ‘vegan’ (Oksanen et al., 2011) to calculate taxa accumulation curves of the two methods over the whole catchment.

#### Alpha diversity

To compare diversity estimates delivered by the two methods, we used the R packages ‘phyloseq’ (McMurdie and Holmes, 2013, version 1.22.3) to calculate richness. With a Pearson’s correlation test we identified if richness measures at sampling sites correlate between the two contemporary methods. We then tested if genus richness of the two methods increased with discharge, using a linear model.

#### Beta diversity

We used Sørensen dissimilarity as a measure of beta-diversity, based on presence/absence data. With the R package ‘betapart’ (Baselga et al., 2017) we calculated Sørensen dissimilarity, which can be further partitioned into nestedness and turnover components, allowing us to distinguish dissimilarity arising through embedment (i.e. loss) or replacement of species (Baselga, 2010, Baselga et al., 2017). To observe how beta diversity measures relate to stream distance, we extracted distances between sites and nodes from GIS data (swisstopo) and added these as edge weights. We then constructed the adjacency matrix representing our fluvial network consisting of edges and vertices with the R package ‘igraph’ (Csardi and Nepusz, 2006, version 1.2.1) to calculate stream distance between flow-connected sampling points. We used linear models to test if distance and the difference in stream order of the compared sites explain patterns in the beta diversity measures. We compared models including a null model, models containing only one of the explanatory variables, or both variables (with and without interaction), and performed model averaging in order to calculate relative variable importance (R package ‘MuMIn’, Barton, 2009, version 1.40.4). We additionally calculated Sørensen dissimilarity and checked for differences among stream orders. First, we tested for heterogeneous variance with a Bartlett test, and if it was significant, we performed a Kruskal-Wallis test to identify whether there was at least one difference in means. If this was true, then we followed a multiple mean comparison post-hoc test on rank sums (R package ‘pgirmess’, Giraudoux, 2018, version 1.6.9).

#### Between-method community dissimilarity

We also calculated Sørensen dissimilarity and its two components, nestedness and turnover, between eDNA and kicknet samples for each individual site to detect discrepancies in community composition between the two contemporary methods.

## Results

### Gamma diversity

At the regional scale (i.e., the whole catchment level), 96 different EPT genera were historically documented. We found 47 EPT genera with our kicknet samples and 42 genera with our eDNA samples, reflecting 46% and 42% of the historically established taxa richness, respectively (Fig. 2, Table S4). A high proportion of these sampled taxa were already present in the historic records (94% and 95% respectively). Thirty-six of these EPT genera were detected with both the kicknet and the eDNA method, reflecting a 62% overlap of the two methods. Both methods also detected a similar additional proportion of taxa found by one method only, but present in the historic dataset (Fig. 2). Finally, we found four genera with eDNA, kicknet sampling, or both approaches that were not previously listed in the historic data. All of these four genera occur at the border of our studied catchment and rare, single appearances within the catchment are possible. Separate taxon accumulation curves (accumulating gamma richness with number of sites included) for the kicknet and the eDNA methods were qualitatively similar (Fig. S2A), based on visual comparison of the curves and their 95% confidence band. This pattern remained similar even if sites were accumulated in the order from the most downstream to the most upstream sites (Fig. S2B).

**Fig. 2:**
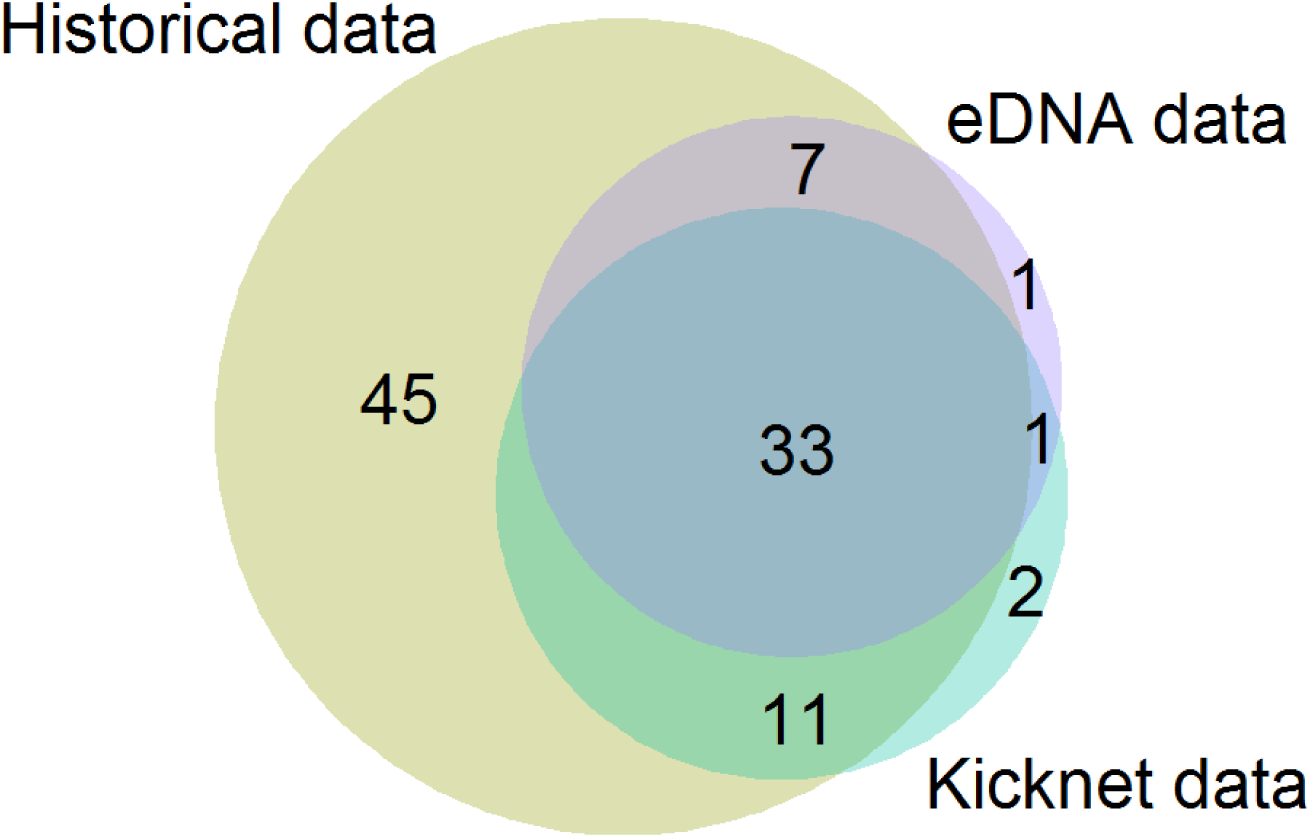
Overlap of EPT genera in the three different datasets over the whole catchment. Bubble size is proportionate to the number of genera detected.

### Alpha diversity

At the local scale, the two contemporary methods revealed similar genus richness (eDNA *M* = 10.22, *SD* = 4.2; kicknet *M* = 10.47, *SD* = 3.1), and the detected richness values were positively correlated (*r*(58) = 0.42, *P* < 0.001; Fig. 3). As eDNA is transported through the river network, we expected a positive dependency of the detected richness of eDNA on increasing discharge (i.e., in more downstream sites), which we do not expect for kicknet samples. Indeed, we found a positive relationship between genus richness and discharge for eDNA (β = 0.537, *t*(58) = 2.848, *p* = 0.006) but not for kicknet (β = 0.073, *t*(58) = 0.495, *p* = 0.62), however, the adjusted R^2^ was relatively low (eDNA R^2^ = 0.101, Fig. 4).

**Fig. 3:**
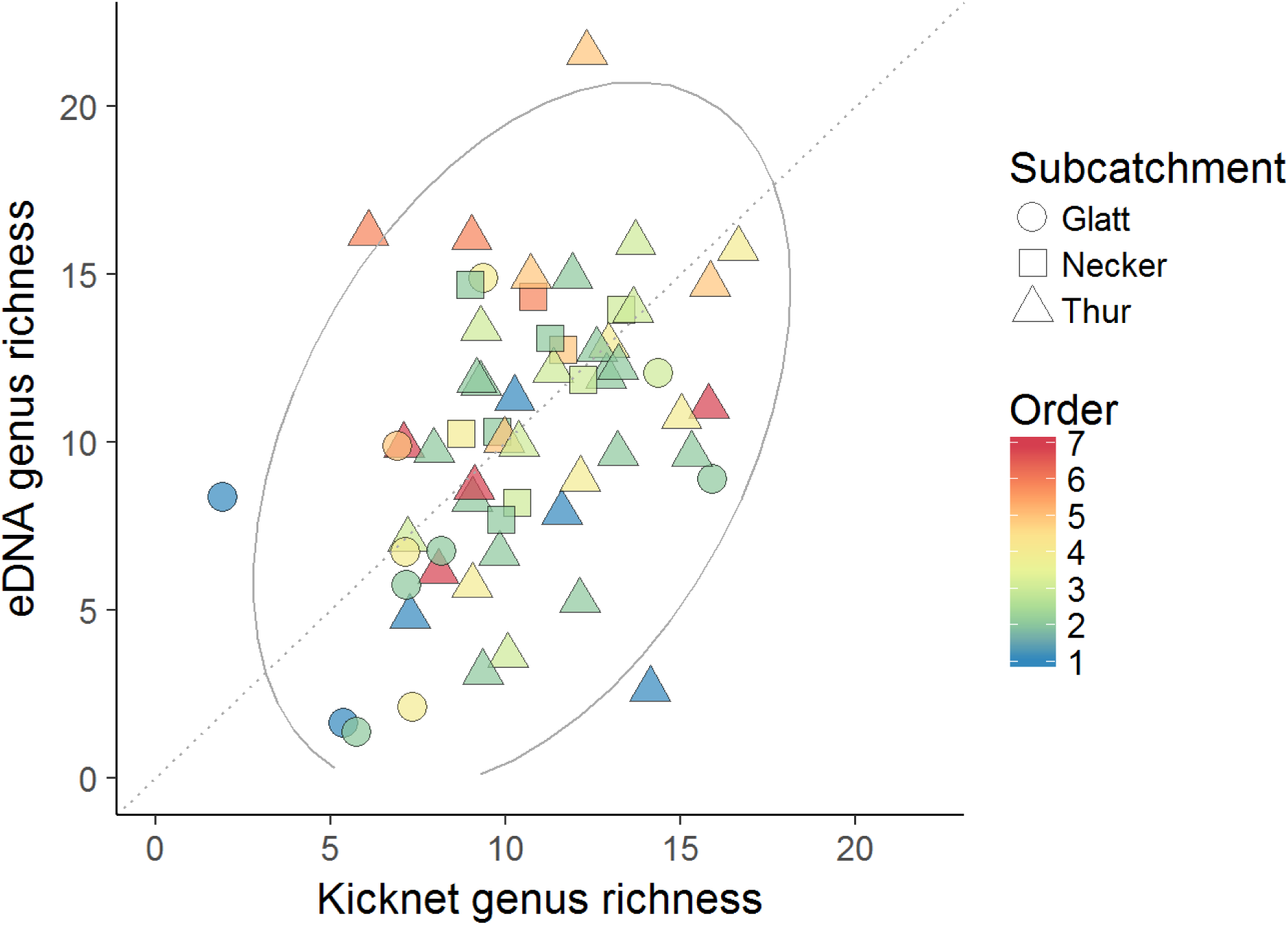
Detected richness of EPT genera comparing kicknet versus eDNA sampling. The grey line indicates the confidence ellipse based on multivariate normal distribution and the dotted line is the 1:1 line. Colors and shapes of the individual data-points are according to stream order and the sampling sites’ subcatchment respectively. Points are minimally jittered due to overlapping cases.

**Fig. 4:**
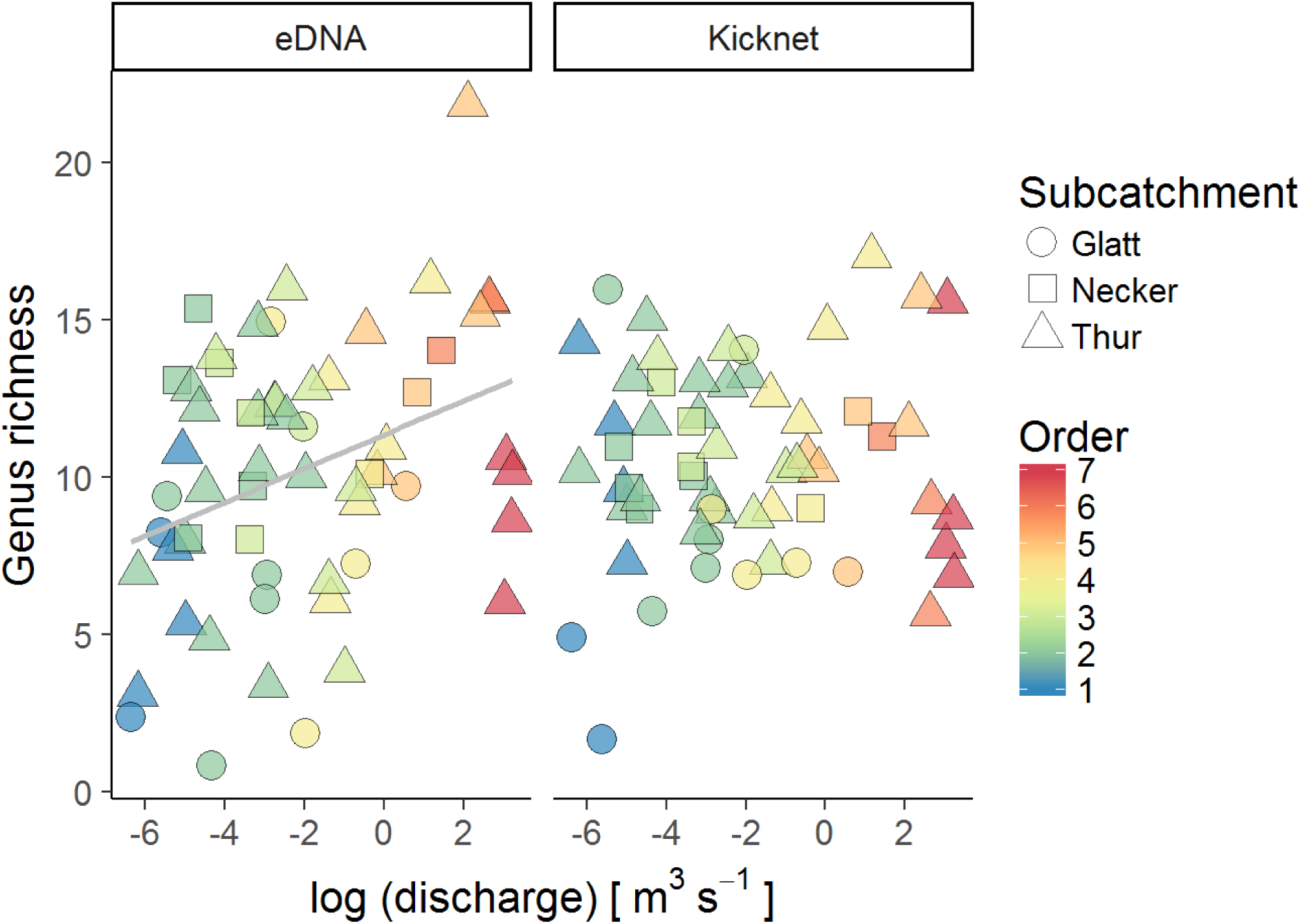
Richness of eDNA and kicknet samples plotted against the logarithmic annual mean discharge. The grey line gives the significant linear regression line for eDNA only due to non-significance for kicknet genus richness. Colors are according to stream order and the shape indicates to what subcatchment the site belongs to. Points are minimally jittered due to overlapping cases.

### Beta diversity

Overall, we found comparable Sørensen dissimilarity between flow-connected sites for eDNA and kicknet. For both methods, the turnover component contributed more to the dissimilarity than nestedness-related components (Fig. 5, see Table S5 for information on detected genera per site), which was more pronounced for eDNA than kicknet. The analysis for the relative importance of variables showed that differences in stream order were generally more important for all beta diversity measures, however, for turnover of both eDNA and kicknet, pairwise distance between sites showed only partially lower importance compared to the differences in stream order (Table S6). Both methods showed significant differences in mean Sørensen dissimilarity among stream orders, and we found significant group differences for the post-hoc mean comparisons in stream orders (Fig. S3, Table S7).

**Fig. 5:**
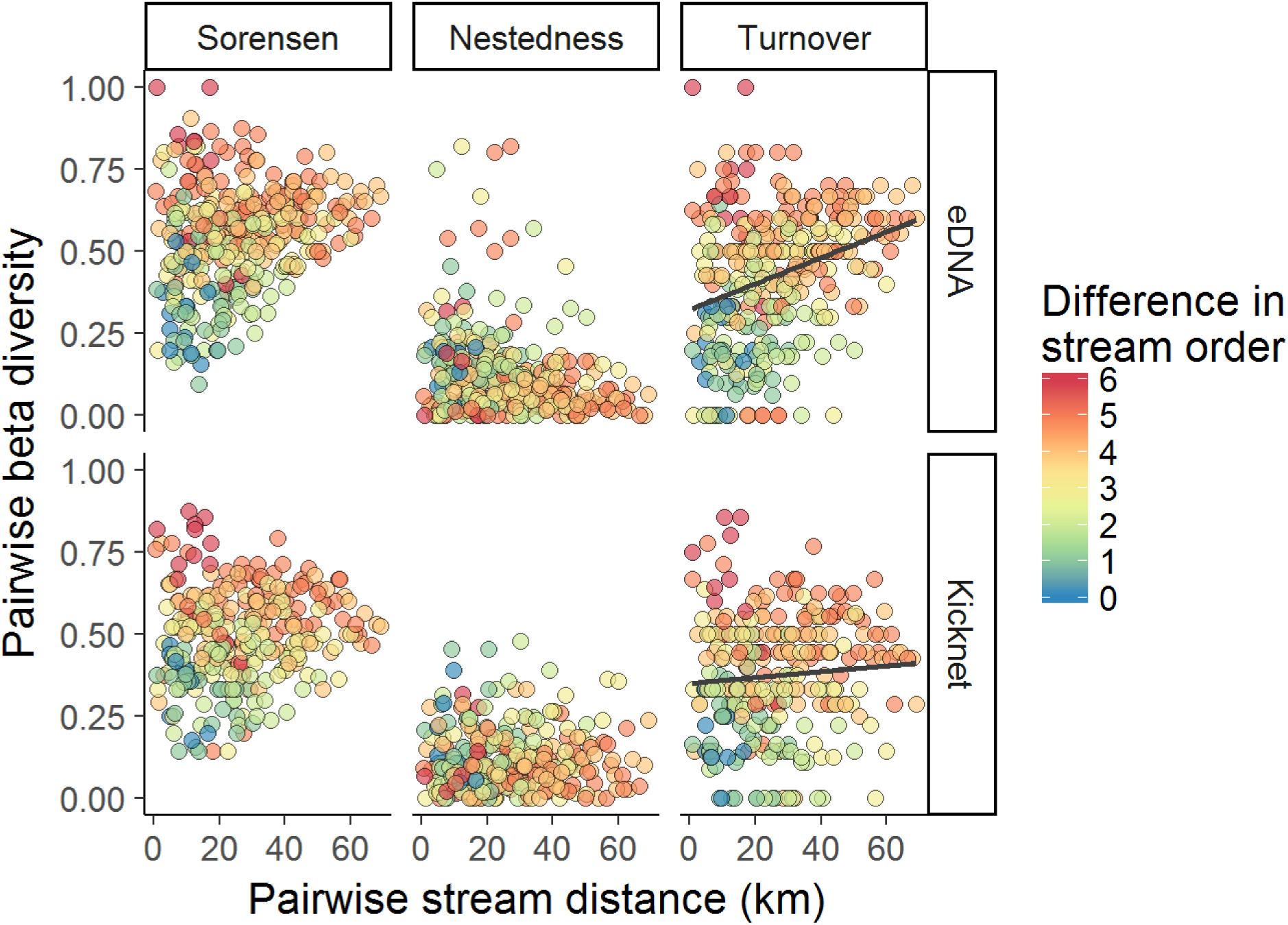
Pairwise beta diversity against pairwise stream distance for flow connected sites only. Beta diversity is calculated based on Sorensen dissimilarity and is split into the two components of nestedness and turnover. The color gradient indicates the difference in stream orders of the pairwise compared sites and solid lines indicate the regression line in cases where stream distance showed relative importance too.

### Between-method community dissimilarity

We found an intermediate discrepancy in the community compositions described by the two contemporary methods (*M* = 0.48, *SD* = 0.14, Fig. S4), and partitioning this difference into nestedness and turnover components indicates that turnover contributes more to the differences of detected EPT genera.

## Discussion

We compared different measures of biodiversity of EPT sampled by kicknet versus by eDNA, using a spatially structured approach and covering a major river network. Using a unique historically assembled overview of cumulative, “true” gamma diversity within the study region, we were able to put our contemporary samples in a historic context. We found a quantitatively similar overlap between each contemporary approach and historic gamma diversity, in accordance with other studies comparing eDNA with long-term data in freshwater systems (e.g., Hänfling et al., 2016, Valentini et al., 2016). Given that the historic data set covers several decades (1981–2016), we do not expect that all taxa are still present in the catchment at the time of this study (2016) due to changes in distribution or local extinctions, and we would also expect some new taxa to appear. Thus, the observed 42% overlap between a single snapshot sampling campaign and the gamma diversity obtained by cumulative sampling efforts over almost four decades is relatively high. Surprisingly, taxon accumulation curves for eDNA and kicknet sampling were not significantly different, although we expected that downstream transport of DNA would contribute to a faster increase and saturation of genera for eDNA compared to kicknet sampling. Our approach suggests that a single eDNA sampling campaign may cover large parts of historically detected gamma diversity. This indicates that eDNA could be used as a new tool for rapid network-level richness analyses and biodiversity assessments. Such systematic BioBlitz sampling campaigns (Lundmark, 2003, Laforest et al., 2013) across hitherto understudied complete river catchments may be a promising avenue, since minimally trained people without any taxonomic expertise can collect great parts of regional or even landscape richness in a rapid time frame with this method.

We also found a reasonable congruency in local alpha diversity of the two contemporary methods and identified a new dependency of eDNA estimates on discharge level. Overall, local richness estimates of eDNA and kicknet sampling are highly comparable. This comparability may be strengthened by our highly stringent inclusion criteria for eDNA estimates which are more conservative than those used in previous studies that detected higher richness with eDNA compared to traditional methods (e.g., Deiner et al., 2016, Valentini et al., 2016). In addition, we used a barcoding primer targeting a broad taxonomic range, which may also have a lower detection rate for specific taxonomic groups. In near future, the forthcoming design and use of EPT specific primers or the completion of EPT sequence references should reduce the current drawbacks of eDNA methods regarding taxonomic identification. For eDNA we found a positive relationship between richness and discharge, which was not the case for kicknet data and could be attributed to eDNA integrating biodiversity detection across space due to downstream transport (Deiner et al., 2016, Li et al., 2018, Pont et al., 2018). Alternatively, richness indeed increased with discharge (i.e. downstream), but the slightly sub-optimal sampling period (summer) and the simplified kicknet protocol we used was an inappropriate technique for sampling larger streams and thus obscured the relationship. Both aspects reduce detection of genera with the traditional methods, as some species might be missed due to reduced sampling effort or immature larval stages hampering identification. Surprisingly, the 7^th^ stream order sites diverge from this pattern, potentially because we could only access part of these wide river cross-sections, and sampling was restricted to the river edge where less mixing occurs, where more extensive sampling may be needed with eDNA (Bylemans et al., 2018). Also, recent studies indicate that eDNA is not evenly distributed in the water column (Macher and Leese, 2017), and it is still unclear how spatial variance in structures, such as riffles and ponds, are affecting the mixing of the water column and thus the equal detection of eDNA along the water column. We speculate that only in smaller streams (1^st^ to 5^th^ order) one or two samples from the edge or in the center adequately reflect the eDNA distribution across the river transect, while in larger rivers multiple samples across the cross-section may be recommended.

We found that pairwise beta diversity (Sørensen dissimilarity) at the local scale was similarly assessed by the two methods, which is congruent to findings by Li et al. (2018). Despite differences among sites in Sørensen dissimilarity, the turnover component (i.e., species replacement) contributes more to the dissimilarity than nestedness for both methods but is even more pronounced for eDNA than kicknet. Turnover indicates that species are replaced between the sites and implies that transportation of eDNA is not the main mechanisms driving differences, otherwise nestedness would be expected to be stronger. Barnes & Turner (2016) presented many processes (e.g., degradation, re-suspension, or fragmentation) that influence eDNA in the environment and therefore challenge our mechanistic understanding. Thus, it remains unclear whether the detected differences stem from ecological or methodological variation.

Studies on diversity patterns have long focused on a linear view of streams. But it is now increasingly acknowledged that the underlying network structure plays a significant role in shaping species distributions (Carrara et al., 2012, Harvey and Altermatt, 2019, Harvey et al., 2018, Holyoak et al., 2005, Seymour et al., 2015, Tonkin et al., 2018). While experimental lab studies and field surveys have found general patterns of diversity distribution in river landscapes, we do not know if eDNA will lead to similar findings or not. Multiple studies showed that there is higher beta diversity among headwaters compared to downstream reaches (Altermatt et al., 2013, Carrara et al., 2012, Finn et al., 2011). When comparing beta diversity within stream orders we detect differences between groups of stream orders for both methods. These differences are mainly between beta diversity of large stream orders and small stream orders, as expected by theory.

Overall, for local composition, we see some discrepancy of the two methods for community composition at specific sites. This discrepancy is driven by turnover, indicating that detection of different species with the two methods differs, and is not due to the integration of eDNA over distance. We see several mechanisms which could cause this difference between the two methods. First, bias introduced in the lab process may have hampered the detection of species that were actually present (e.g., through primer bias (Elbrecht and Leese, 2017), PCR stochasticity (Leray and Knowlton, 2017), sequencing depth, etc.). The design of specific EPT primers may improve comparison and deliver even a better overlap than a universal eukaryotic primer as we used in this study. Second, DNA shedding rates, densities, and activity can differ between genera (Bylemans et al., 2017, de Souza et al., 2016, Sassoubre et al., 2016) and affect the eDNA detection. Also, it remains unknown how habitat preferences of different genera affect detection due to limited mixing or flow of the preferred habitat. Third, it is possible that DNA got re-suspended from sediments through the mixing of the water column or other disturbances (Jerde et al., 2016, Shogren et al., 2017, Shogren et al., 2016), resulting in a signal of locally extinct taxa. However, we have no indication from the historic data that genera detected only with eDNA have been absent in the catchment for a longer time.

## Conclusion

The distribution of biodiversity and indicator species is of interest to many fields in ecology, from basic research and theory to applied projects of biodiversity conservation and ecosystem assessments. Our work identifies novel opportunities of eDNA as a reliable tool to detect biodiversity patterns, similar to traditional kicknet sampling in riverine networks. Our findings show high robustness of the method, allowing its use for rapid richness analyses and offering the potential to do quick assessments of biodiversity with untrained collectors for BioBlitz campaigns or citizen science projects. However, if the goal is to extend previous biomonitoring or biodiversity datasets with eDNA sampling, the spatial scale must be considered when designing sampling schemes to detect taxon richness.

## Authors’ contributions

EM, CJL, RA, EF, IG, EH, JCW and FA conceived the ideas and designed methodology; EM, CJL, RW, RA, EF, IG, EH, SH, JCW, and FA collected the data; EM, JCL, RW, and JCW analysed the data; EM and FA led the writing of the manuscript. All authors contributed critically to the drafts and gave final approval for publication.

## Acknowledgements

Data analyzed in this paper were generated in collaboration with the Genetic Diversity Centre (GDC), ETH Zurich. We would like to thank Simon Flückiger and Sereina Gut for help in the lab and Rosi Sieber for support with the GIS data. Special thanks go to Rosetta Blackman, Kristy Deiner, and Hanna Hartikainen for inputs on the manuscript. We thank the Centre Suisse de Cartographie de la Faune (CSCF)/InfoFauna for the historic data on EPT in the Thur catchment. Funding is from the Swiss National Science Foundation Grants No PP00P3_179089 and 31003A_173074, the Velux Foundation, and the University of Zurich Research Priority Program “*URPP Global Change and Biodiversity”* (to F.A.), and the Forschungskredit grant of the University of Zürich (to IG). This is publication ISEM-YYYY-XXX of the Institut des Sciences de l’Evolution - Montpellier.

## Data accessibility

Sequencing data will be publicly available on Dryad after publication.

## Supporting information

See supplementary file attached to the paper.

