## Supplemental File for "Assessing different components of biodiversity across a river network using eDNA"

### Supplementary file for submission to Journal of Applied Ecology: Assessing different components of biodiversity across a river network using eDNA

Elvira Mächler[1,2*†], Chelsea J. Little[1,2†], Remo Wüthrich[1], Roman Alther[1,2], Emanuel A. Fronhofer[1,2,3], Isabelle Gounand[1,2], Eric Harvey[1,2,4], Samuel Hürlemann[1], Jean-Claude Walser[5], Florian Altermatt[1,2*]

[1] *Eawag: Swiss Federal Institute of Aquatic Science and Technology, Department of Aquatic Ecology, Überlandstrasse 133, CH-8600 Dübendorf, Switzerland.*
[2] *Institute of Evolutionary Biology and Environmental Studies, University of Zurich, Winterthurerstrasse 190, CH-8057 Zürich, Switzerland.*
[3] *ISEM, Univ Montpellier, CNRS, EPHE, IRD, Montpellier, France.*
[4] *Département de Sciences Biologiques, Université de Montréal, Montréal, Canada, H2V 2S9*
[5] *Federal Institute of Technology (ETH), Zürich, Genetic Diversity Centre, CHN E 55 Universitätstrasse 16, 8092 Zürich, Switzerland*

[†] shared first authorship

### Contents


**1 Table S1: Sampling sites** **2**

**2 Detailed information on field filtration, eDNA facilities and material preparation** **4**

**3 Detailed information on eDNA extraction** **4**

**4 Detailed information on library preparation** **5**

**5 Detailed information on dummy sequence for positive control** **6**

**6 Detailed information on statistics** **7**

**7 Table S2: Primer sequences** **8**

**8 Table S3: Number or reads** **9**

**9 Figure S1: Shared ZOTUS within sites** **10**

**10 Table S4: Genera list** **11**

**11 Figure S2: Genera accumulation curves** **15**

**12 Table S5: Detected genera per site** **16**

**13 Table S6: Relative importance** **16**

**14 Figure S3: Beta diversity within stream orders** **17**

**15 Table S7: Statistics to Figure S3** **18**

**16 Figure S4: Beta diversity between sampling methods** **19**

**17 References** **20**

### 1 Table S1: Sampling sites

**Table S1:** Overview of sampling sites in the presented study. Note that site 38 is missing due to inaccessibility during fieldwork.

| Site number | Site name | X coordinate | Y coordinate | Sub-catchment | Stream order | Discharge |
|---|---|---|---|---|---|---|
| 1 | Gertenau | 726'445 | 262'809 | Thur | 1 | 0.007 |
| 2 | Niederuzwil | 728'545 | 256'867 | Thur | 4 | 0.260 |
| 3 | Sagenbach | 724'173 | 248'426 | Thur | 6 | 14.380 |
| 4 | Tannenhof | 725'234 | 257'261 | Thur | 2 | 0.007 |
| 5 | Stocken | 725'111 | 257'884 | Thur | 7 | 21.450 |
| 6 | Rickenbach | 721'110 | 256'854 | Thur | 3 | 0.250 |
| 7 | Aueli | 723'409 | 254'006 | Thur | 7 | 20.543 |
| 8 | Aspis | 723'535 | 253'665 | Thur | 1 | 0.002 |
| 9 | Bruggbach | 720'262 | 252'586 | Thur | 2 | 0.140 |
| 10 | Letten | 733'018 | 259'732 | Thur | 7 | 25.384 |
| 11 | Erlenhof | 733'111 | 258'661 | Thur | 3 | 0.370 |
| 12 | Ruteli | 729'223 | 257'447 | Thur | 7 | 24.724 |
| 13 | Chruxalpen | 731'245 | 257'766 | Thur | 2 | 0.063 |
| 14 | Oberriet | 725'911 | 253'731 | Glatt | 2 | 0.004 |
| 15 | Rotelbach | 740'671 | 256'165 | Glatt | 2 | 0.053 |
| 16 | Riederen | 731'721 | 253'993 | Glatt | 1 | 0.002 |
| 17 | Glatthalde | 733'000 | 252'973 | Glatt | 5 | 1.770 |
| 18 | Botsberg | 730'375 | 252'731 | Glatt | 4 | 0.140 |
| 19 | Burgau | 733'895 | 251'587 | Glatt | 1 | 0.004 |
| 20 | Matt | 732'733 | 251'496 | Glatt | 2 | 0.050 |
| 21 | Oberrindal | 727'885 | 251'736 | Thur | 1 | 0.005 |
| 22 | Muhlau | 723'560 | 252'131 | Thur | 7 | 20.270 |
| 23 | Gahwil | 717'944 | 250'623 | Thur | 2 | 0.054 |
| 24 | Zellersmuehli | 737'293 | 251'374 | Glatt | 4 | 0.490 |
| 25 | Wissbach | 735'628 | 246'601 | Glatt | 4 | 0.058 |
| 26 | Vier Winden | 739'290 | 247'888 | Glatt | 2 | 0.013 |
| 27 | Landersberg | 737'716 | 247'080 | Glatt | 3 | 0.130 |
| 28 | Hengarten | 725'771 | 249'605 | Necker | 2 | 0.036 |
| 29 | Altstig | 727'103 | 248'360 | Necker | 6 | 4.180 |
| 30 | Hoffeld | 731'136 | 247'220 | Necker | 2 | 0.009 |
| 31 | Winklen | 720'873 | 248'407 | Thur | 5 | 0.630 |
| 32 | Muhlruti | 716'420 | 247'168 | Thur | 2 | 0.002 |
| 33 | Bodmen | 717'992 | 245'312 | Thur | 3 | 0.086 |
| 34 | Bitzi | 722'422 | 247'463 | Thur | 1 | 0.006 |
| 35 | Neudietfurt | 724'130 | 245'284 | Thur | 6 | 14.020 |
| 36 | Ritzentaa | 723'525 | 245'098 | Thur | 5 | 0.840 |
| 37 | Buech | 728'234 | 246'422 | Necker | 3 | 0.016 |
| 39 | Gluris | 729'191 | 239'587 | Necker | 3 | 0.034 |
| 40 | Peterzell | 731'142 | 242'155 | Necker | 5 | 2.270 |
| 41 | Eggliwald | 721'123 | 241'713 | Thur | 2 | 0.012 |
| 42 | Stalden | 727'096 | 240'815 | Thur | 3 | 0.065 |
| 43 | Blumenstein | 735'999 | 241'461 | Necker | 2 | 0.007 |
| 44 | Hemberg | 732'060 | 238'913 | Necker | 2 | 0.006 |

- continued on next page -

**Table S1 – continued from previous page**

| Site number | Site name | X coordinate | Y coordinate | Sub-catchment | Stream order | Discharge |
|---|---|---|---|---|---|---|
| 45 | Vordernecker | 733'487 | 237'763 | Necker | 4 | 0.690 |
| 46 | Wattwil | 725'591 | 238'615 | Thur | 5 | 11.160 |
| 47 | Schonenberg | 723'135 | 236'789 | Thur | 2 | 0.011 |
| 48 | Gstaltlig | 728'504 | 236'720 | Thur | 2 | 0.042 |
| 49 | Ricken | 721'544 | 236'108 | Thur | 4 | 0.250 |
| 50 | St.Johann | 732'240 | 232'834 | Thur | 5 | 8.130 |
| 51 | Nestel | 726'804 | 232'818 | Thur | 2 | 0.086 |
| 52 | Zuu | 728'712 | 230'912 | Thur | 4 | 0.540 |
| 53 | Oberwald | 740'357 | 235'200 | Thur | 2 | 0.008 |
| 54 | Rietbad | 736'033 | 234'193 | Thur | 4 | 1.050 |
| 55 | Laui | 742'285 | 231'249 | Thur | 3 | 0.490 |
| 56 | Buechel | 735'257 | 231'681 | Thur | 2 | 0.042 |
| 57 | Sulzbach | 732'740 | 228'431 | Thur | 3 | 0.165 |
| 58 | Seluner | 736'494 | 226'521 | Thur | 2 | 0.010 |
| 59 | Wildhaus | 745'203 | 229'087 | Thur | 3 | 0.015 |
| 60 | Starkenbach | 737'954 | 227'739 | Thur | 4 | 3.180 |
| 61 | Schwendi | 742'567 | 228'161 | Thur | 2 | 0.043 |
| 62 | Tufi | 735'869 | 244'364 | Necker | 3 | 0.034 |

#### 2 Detailed information on field filtration, eDNA facilities and material preparation

After finishing the filtration, we rolled the filters with the help of tweezers, put them in individual 1.5 mL tubes and stored them in a Styrofoam box equipped with cooling elements until we came back from the field (no longer than 9 h). Thereafter, they were stored at -20 °C until further processing. Tweezers were cleaned with 2.5% bleach, rinsed with 100% molecular grade EtOH and dried with household paper at beginning of a new site, but not in-between replicates of one site. On each field day, we performed a filter control (FC) that consisted of 250 mL of nanopore water previously treated with UVC light and sealed in the clean lab facilities. The filter control was replicated four times and was filtered in the field before any sampling site was visited in order to check if the reusable material was clean. In total, we generated 11 filter controls.

All eDNA sampling material preparation and extractions were performed in a lab facility dedicated to perform sensitive eDNA work, especially equipped with a positive air pressure and separated from all PCR environments. Reusable materials (filter housings (Swinnex, EMD Millipore Co., Billerica, Massachusetts), sterile syringes) were cleaned in 2.5% bleach, rinsed with ddH2O and dried inside the clean lab facility. After drying, the material was treated with UVC light together with the glass fibre filters (GF/F, Whatman International Ltd., Maidstone, U.K.) and 1.5 mL tubes. Afterwards, the filters were put into the filter housings and we prepared for each sampling site a zip lock bag that contained a syringe, four filter housings, and four 1.5 mL tubes. These bags were only opened at a specific sampling site.

#### 3 Detailed information on eDNA extraction

We randomized the samples for the extraction. 12 samples were extracted together in one batch and each of the batches contained at least one filter control. Per extraction day we did maximally 3 batches (i.e., 36 samples) whereof one was an extraction control (EC), in total this resulted in 8 extraction controls. The extraction control consisted of a GF/F filter that was treated with UVC light beforehand. We used the DNeasy Blood and Tissue kit (Qiagen GmbH, Hilden, Germany) following the spin column protocol for animal tissues besides the following changes: First, we doubled the reaction volumes until the loading of the mixture on the spin column (step 4 in the protocol). Second, we eluted twice with 37.5 µL and pooled the two elutions together, resulting in 75 µL of eDNA per sample. Each sample was then cleaned up with the One Step PCR inhibitor removal kit (Zymo, Research, Irvine, California).

### 4 Detailed information on library preparation

We used a two-step protocol to prepare the library for each sample. The first PCR was performed with primers that contained a nexttera adaptor, a sequencing site, heterogeneity spacer and the primer sequence (see Table S2). The first PCR consisted of 1X buffer I (3 µL), 1.25 U AmpliTaq Gold (0.25 µL), 0.1 mg/mL BSA (0.3 µL), 0.2 mM dNTPs (0.6 µL), 1.0 mM MgCl2 (1.2 µL), 0.5 µM forward primer mix (1.5 µL), 0.5 µM reverse primer mix (1.5 µL) and 5 µL eDNA template in a total volume of 30 µL. The samples were randomized on three PCR plates for the library preparation. On each of the PCR plates, we implemented one negative (NC) and one positive PCR control (PC), resulting in a total of 3 controls of each. The negative control consisted of 5 µL sigma water and the positive control consisted of 4 µL eDNA sample and 1 µL(0.01 ng/µL) dummy DNA (See below: 5 Detailed information on dummy sequence for positive control). The PCR regime consisted of 95 °C for ten minutes, followed by 44 cycles of denaturation at 95 °C for 15 seconds, annealing at 62 °C for 30 seconds and elongation at 72 °C for 45 seconds, ending the PCR with a final hold of 72 °C for 5 minutes. The amplification success was verified with the QiAxcel Screening Cartridge by using the AL420 method (Qiagen, Hilden, Germany). If the amplicon did not successfully amplify, we tested a 1:10 dilution of the eDNA sample (5 µL) and checked again for success. We selected three sampling replicates per site for the further library preparation based on the preliminary test. For each sample, we performed 3 PCR replicates (with either 5 µL of eDNA or 5 µL of 1:10 diluted eDNA, depending on the amplification success, see Table Sx). We pooled the three replicates per site and took 30 µL to clean with SPRI beads (ratio 0.8x) following the manufacturer's protocol. The clean product was eluted in 20 µL sigma water and we were able to recover 18 µL of the cleaned product.

For the second PCR, we used the Nextera XT Index kit v2 to index each sample. The reaction consisted of 1X KAPA HiFi (25 µL), 5 µL Index S, 5 µL Index N and 15 µL cleaned template in a total of 50 µL per reaction. The PCR regime consisted of 95 °C for 3 minutes, followed by 10 cycles of denaturation at 95 °C for 30 seconds, annealing at 55 °C for 30 seconds and elongation at 72 °C for 30 seconds, ending the PCR with a final hold of 72 °C for 5 minutes. We took 25 µL of the index reaction and cleaned it with SPRI beads, ratio 0.8 x (20 µL) following the manufacturer's protocol. The clean index reactions were resolved in 20 µL sigma water and we recovered 18 µL of the clean indexed product.

We quantified each sample with the Spark 10M Multimode Microplate Reader (Tecan® Group Ltd., Männedorf, Switzerland) by using the Qubit dsDNA BR assay. Measurements were done in duplicates on a 384-well plate with an assay volume of 50 µL and 2 µL of cleaned index reaction. We calculated the concentration of each sample based on an average library size of 535 bp. In order to create feasible pipetting volumes, we normalized the samples into 5 different pools depending on their concentration by using a liquid handling station (BRAND, Wertheim, Germany). We then measured the concentrations of each generated pool with the Qubit dsDNA BR assay. Based on the measured concentration, we then pooled the five pools equimolar into a final pool. The final pool, containing all samples, was cleaned with SPRI beads (0.8x ratio) and a final check was done on the Agilent 2200 TapeStation High Sensitivity D1000 assay (Agilent Technologies, California, USA). All controls (FC, EC, PC, NC) were run along to the samples and were pooled according to their concentrations. In case the negative controls were blank (i.e. no concentration could be estimated), they were added to the second lowest concentrated pool with 10 µL.

### 5 Detailed information on dummy sequence for positive control

The dummy DNA consisted of a randomly generated dsDNA sequence that was 313 bp in length and has 38% GC-content, matching the expected amplified amplicon from previous studies (Lim et al. 2016). The sequence ordered is the following (bold letters indicate primer binding sites) and was assigned to ZOTU7:

**GGAACAGGTTGAACTGTATATCCCCC**ATCAACCTAGTTACGAAGAGCTATAGATCATAT
AATCCTTAAGTGGAATGTTAATGTGAGTTCAATATGATACACGCCACAGACTCATGTATGTG
GATCGGAAGCCAGCTGTTTCCGACCTCGGAGCCGAGAGTGGTTTCTGAATTACACATGTAA
GATAAAATCATTAAAGGTACTAACTCACGAAACCTCAGGATATGCGTGGTTTGCTGAGATTT
CTATTTTCTCGTTCTTGATTTAACCACGTAAAATGTGTGAAAACTAAAGGTTCTAGCATTTC
TAAGGATCACTACGCCTAACGTCTCACTTTATCTTAAATT**TGATTTTTTGGTCACCCTGA
AGTTTA**

### 6 Detailed information on statistics

First, we verified that the two Illumina runs could be combined. Therefore, we transformed the abundance data into relative abundance data and then calculated the Bray-Curtis distance between the two runs using the R package 'vegan' (Oksanen et al. 2010, version 2.4-6). We analyzed the variance among runs and sites by applying an Adonis test to the Bray-Curtis distance matrices. Further, we tested for the multivariate homogeneity of group dispersions using the two Illumina runs as a grouping factor. As both tests were not significant (i.e. there was no difference in the means or dispersion between the two runs), we were able to combine the data. As a stringency measure, we only counted ZOTUS that were present in at least two of the three eDNA replicates at a specific site. We did not apply an abundance threshold used in other eDNA studies (e.g., Macher et al. 2017), as this did not improve the percentage of shared ZOTUS among site replicates (Fig. S1). Next, we removed all ZOTUs that were present in the negative controls from our field samples and selected only ZOTUs that were assigned either to an Ephemeroptera, Plecoptera or Trichoptera order. We performed all analyses at the genus level, as both eDNA and kicknet datasets had incomplete individual/sequence assignment at the species level due to cryptic species, immature larvae, or sequences not being assigned to that taxonomic resolution .

#### 7 Table S2: Primer sequences

**Table S2:** Forward and reverse primer sequence for the first, tailed PCR reactions. Bold letters indicated the adaptor specific tail and italic letters represent the heterogeneity spacer.

| Primer name | Primer sequence |
|---|---|
| mlCOIintF_FS0 | **TCGTCGGCAGCGTCAGATGTGTATAAGAGACAG**GGWACWGGWTGAACWGTWTAYCCYCC |
| mlCOIintF_FS1 | **TCGTCGGCAGCGTCAGATGTGTATAAGAGACAG***G*GGWACWGGWTGAACWGTWTAYCCYCC |
| mlCOIintF_FS3 | **TCGTCGGCAGCGTCAGATGTGTATAAGAGACAG***TAG*GGWACWGGWTGAACWGTWTAYCCYCC |
| jgHCO2198_FS0 | **GTCTCGTGGGCTCGGAGATGTGTATAAGAGACAG**TAIACYTCIGGRTGICCRAARAAYCA |
| jgHCO2198_FS1 | **GTCTCGTGGGCTCGGAGATGTGTATAAGAGACAG***T*TAIACYTCIGGRTGICCRAARAAYCA |
| jgHCO2198_FS2 | **GTCTCGTGGGCTCGGAGATGTGTATAAGAGACAG***TA*TAIACYTCIGGRTGICCRAARAAYCA |

#### 8 Table S3: Number or reads

**Table S3:** Number of ZOTUs and numbers of reads remaining in the data set after applying specific filtering steps.

| Applied step | Number of ZOTUs | Number of reads |
|---|---|---|
| Raw data | 11,313 | 27,354,262 |
| Presence in minimum two replicated samples | 6,036 | 24,070,851 |
| Removal of ZOTUS present in negative controls | 5,714 | 9,874,010 |
| Assigned to EPT order | 131 | 912,015 |

#### 9 Figure S1: Shared ZOTUS within sites

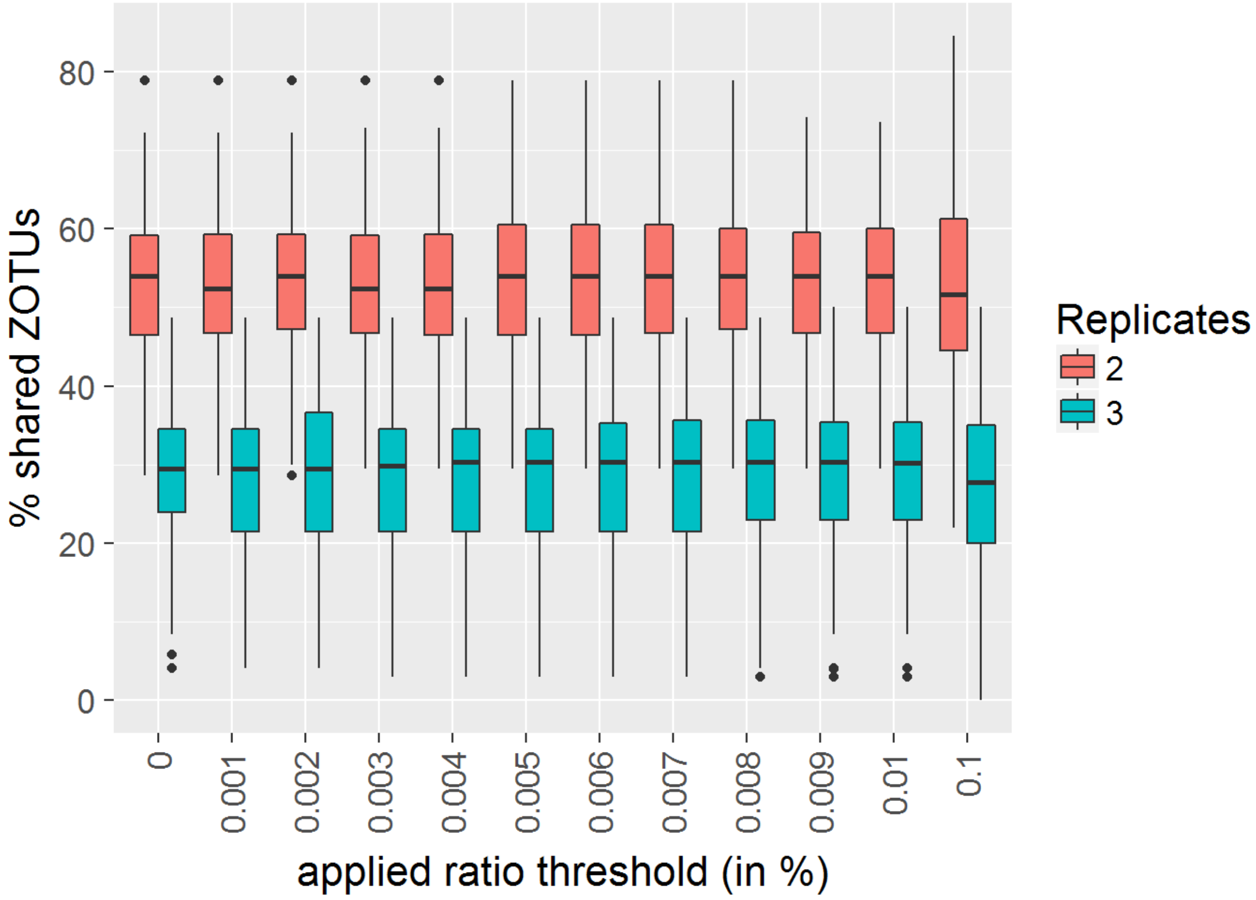


**Fig. S1:** Percent shared ZOTUS for two or three replicates within a site versus different applied ratio thresholds.

### 10 Table S4: Genera list

**Table S4:** List of genera present in Historic data set or detected with eDNA and kicknet. Presences is indicated in green and absence is indicated in red. E stands for Ephemeroptera, P for Plecoptera and T for Trichoptera.

| Genus | Order | Historic | eDNA | Kicknet |
|---|---|---|---|---|
| Adicella | T | green | red | red |
| Agapetus | T | green | red | red |
| Agraylea | T | green | red | red |
| Agrypnia | T | green | red | red |
| Alainites | E | green | green | green |
| Allogamus | T | green | green | green |
| Allotrichia | T | red | green | red |
| Ameletus | E | green | red | red |
| Amphinemura | P | green | green | green |
| Anabolia | T | green | red | red |
| Annitella | T | green | green | red |
| Athripsodes | T | green | green | green |
| Baetis | E | green | green | green |
| Beraea | T | green | red | red |
| Brachycentrus | T | green | red | red |
| Brachyptera | P | green | red | red |
| Caenis | E | green | green | green |
| Capnia | P | green | red | red |
| Capnioneura | P | green | red | red |
| Centroptilum | E | green | red | green |
| Ceraclea | T | green | red | red |
| Chaetopteryx | T | green | red | green |
| Chloroperla | P | green | green | green |
| Cloeon | E | green | green | green |
| Crunoecia | T | green | green | green |
| Cyrnus | T | green | red | red |
| Dictyogenus | P | red | red | green |

- continued on next page -

**Table S4 – continued from previous page**

| Genus | Order | Historic | eDNA | Kicknet |
|---|---|---|---|---|
| Dinocras | P | green | green | green |
| Drusus | T | green | green | green |
| Ecclisopteryx | T | green | red | red |
| Ecdyonurus | E | green | green | green |
| Ecnomus | T | green | red | red |
| Electrogena | E | green | green | green |
| Enoicyla | T | green | red | red |
| Epeorus | E | green | green | green |
| Ephemera | E | green | green | green |
| Ephemerella | E | green | red | red |
| Ernodes | T | green | red | red |
| Erotesis | T | green | red | red |
| Glossosoma | T | green | green | red |
| Glyphotaelius | T | green | red | red |
| Goera | T | red | red | green |
| Habroleptoides | E | green | green | green |
| Habrophlebia | E | green | red | green |
| Halesus | T | green | green | red |
| Heptagenia | E | green | red | red |
| Hydatophylax | T | green | red | red |
| Hydropsyche | T | green | green | green |
| Hydroptila | T | green | green | green |
| Isoperla | P | green | green | green |
| Lepidostoma | T | green | red | green |
| Leuctra | P | green | green | green |
| Limnephilus | T | green | green | green |
| Lithax | T | green | red | red |
| Lype | T | green | red | green |
| Melampophylax | T | green | green | green |
| Mesophylax | T | green | red | red |
| Metanoea | T | green | red | green |

- continued on next page -

**Table S4 – continued from previous page**

| Genus | Order | Historic | eDNA | Kicknet |
|---|---|---|---|---|
| Micrasema | T | | | |
| Micropterna | T | | | |
| Molanna | T | | | |
| Mystacides | T | | | |
| Nemotaulius | T | | | |
| Nemoura | P | | | |
| Nemurella | P | | | |
| Nigrobaetis | E | | | |
| Odontocerum | T | | | |
| Oecetis | T | | | |
| Oligotricha | T | | | |
| Oxyethira | T | | | |
| Parachiona | T | | | |
| Paraleptophlebia | T | | | |
| Perla | P | | | |
| Perlodes | P | | | |
| Philopotamus | T | | | |
| Phryganea | T | | | |
| Plectrocnemia | T | | | |
| Polycentropus | T | | | |
| Potamophylax | T | | | |
| Protonemura | P | | | |
| Psychomyia | T | | | |
| Ptilocolepus | T | | | |
| Rhabdiopteryx | P | | | |
| Rhadicoleptus | T | | | |
| Rhithrogena | E | | | |
| Rhyacophila | T | | | |
| Sericostoma | T | | | |
| Serratella | E | | | |
| Setodes | T | | | |

- continued on next page -

**Table S4 – continued from previous page**

| Genus | Order | Historic | eDNA | Kicknet |
|---|---|---|---|---|
| Silo | T | green | red | red |
| Siphlonurus | E | green | green | green |
| Siphonoperla | P | green | red | red |
| Stactobia | T | green | red | red |
| Stenophylax | T | green | red | red |
| Synagapetus | T | green | green | red |
| Taeniopteryx | P | green | red | red |
| Tinodes | T | green | red | green |
| Tricholeiochiton | T | green | red | red |
| Wormaldia | T | green | green | green |
| Zwicknia | P | green | green | red |

#### 11 Figure S2: Genera accumulation curves

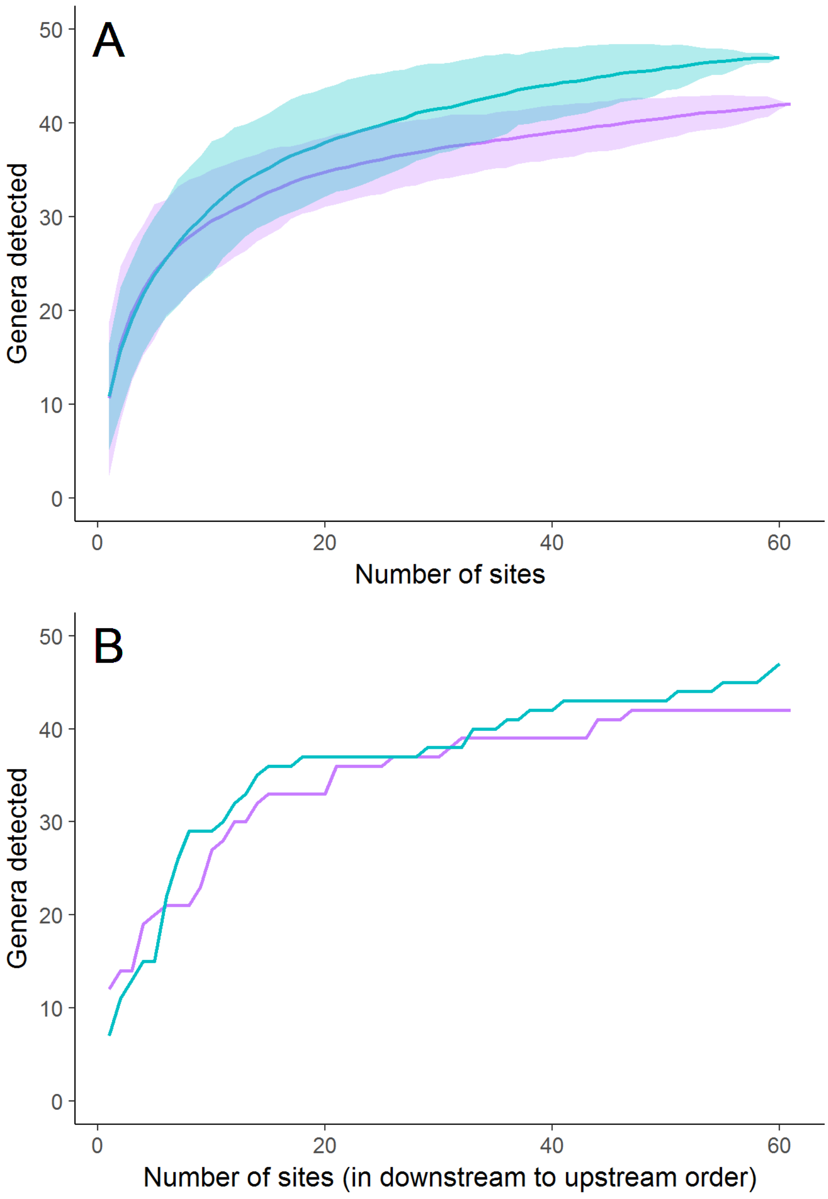


**Fig. S2:** Genera accumulation curves of the two sampling methods (blue Kicknet, purple eDNA). Upper panel **(A)** shows accumulation curve for random sampling and the ribbon indicates the 95% confidence interval. Lower panel **(B)** shows accumulation ordered from the most downsteam site to the most upstream site.

### 12 Table S5: Detected genera per site

See separate excel file containing Table S5.

**Table S5:** Detected Genus per site and method.

### 13 Table S6: Relative importance

**Table S6:** Results table of relative variable importance for each individual dissimilarity measure. Number of models indicates in how many models the variable was used.

| Method | Dissimilarity measure | Difference in stream order | Distance | Interaction |
|---|---|---|---|---|
| eDNA | Sorensen | 1.0 | 0.5 | 0.2 |
| | Nestedness | 0.99 | 0.67 | 0.31 |
| | Turnover | 1.0 | 0.99 | 0.27 |
| Kicknet | Sorensen | 1.0 | 0.55 | 0.16 |
| | Nestedness | 1.0 | 0.88 | 0.7 |
| | Turnover | 1.0 | 0.92 | 0.63 |
| Number of models | | 3 | 3 | 1 |

#### 14 Figure S3: Beta diversity within stream orders

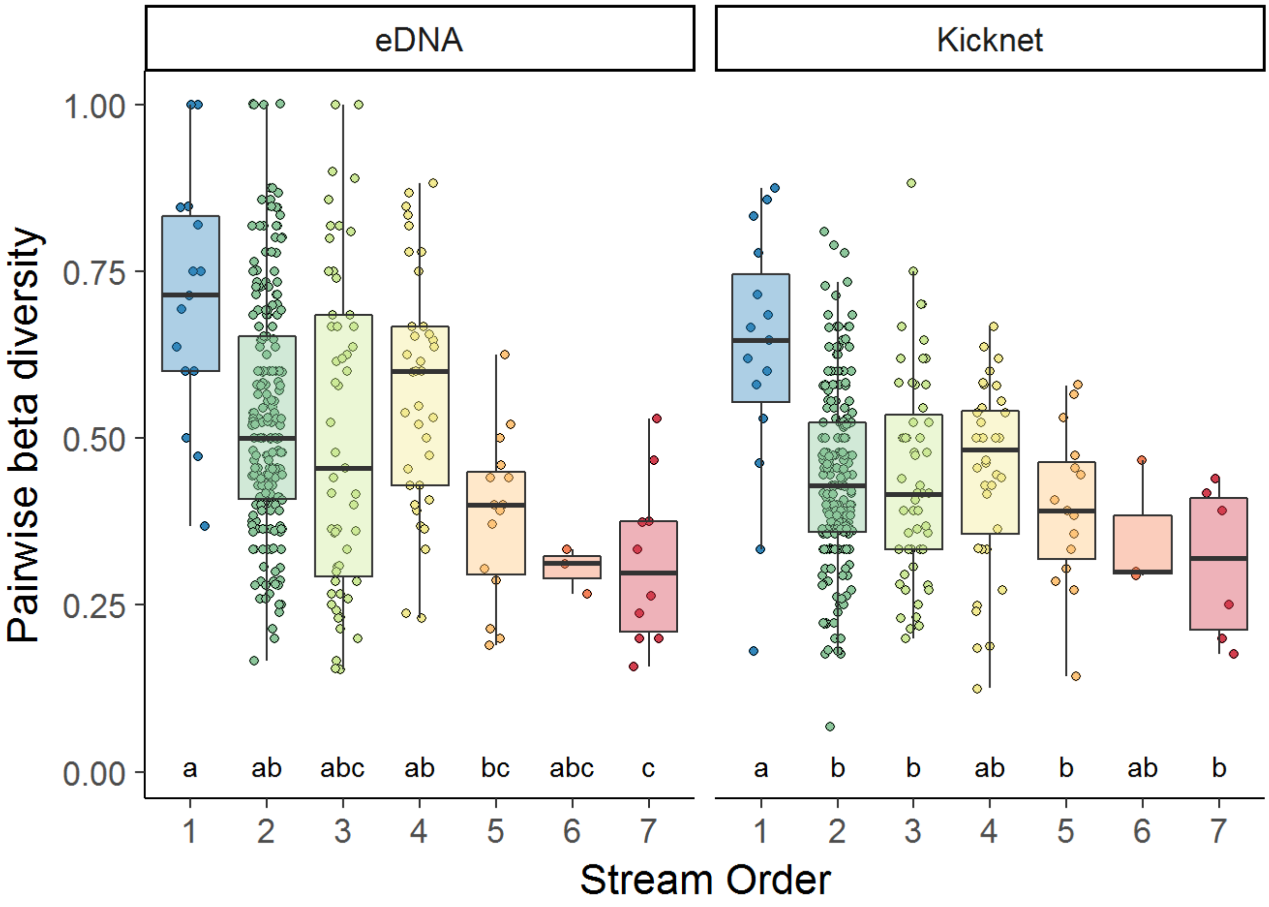


**Fig. S3:** Pairwise beta diversity (Sorensen dissimilarity) within stream orders. Stream orders with the same letter are not significantly different according to multiple mean comparison test after Kruskal-Wallis.

#### 15 Table S7: Statistics to Figure S3

**Table S7:** Results of Kruskal-Wallis test on ranks and the following multiple mean comparisons post-hoc tests based on rank sums. Chi-Squared, degrees of freedom (DF), number of data points (n) and P-values are given. Stream orders with the same letter are not significantly different according to multiple comparison post-hoc test after Kruskal-Wallis (p- value = 0.01).

| Kruskal-Wallis test on ranks | | | | | Multiple mean comparisons post-hoc tests | | | | | | |
|---|---|---|---|---|---|---|---|---|---|---|---|
| **Method** | **Chi-Squared** | **DF** | **n** | **P-value** | **1** | **2** | **3** | **4** | **5** | **6** | **7** |
| eDNA | 40.6 | 6 | 688 | <0.001 | ab | a | ab | b | ac | ac | c |
| Kicknet | 23.37 | 6 | 680 | <0.001 | a | b | b | ab | b | ab | b |

#### 16 Figure S4: Beta diversity between sampling methods

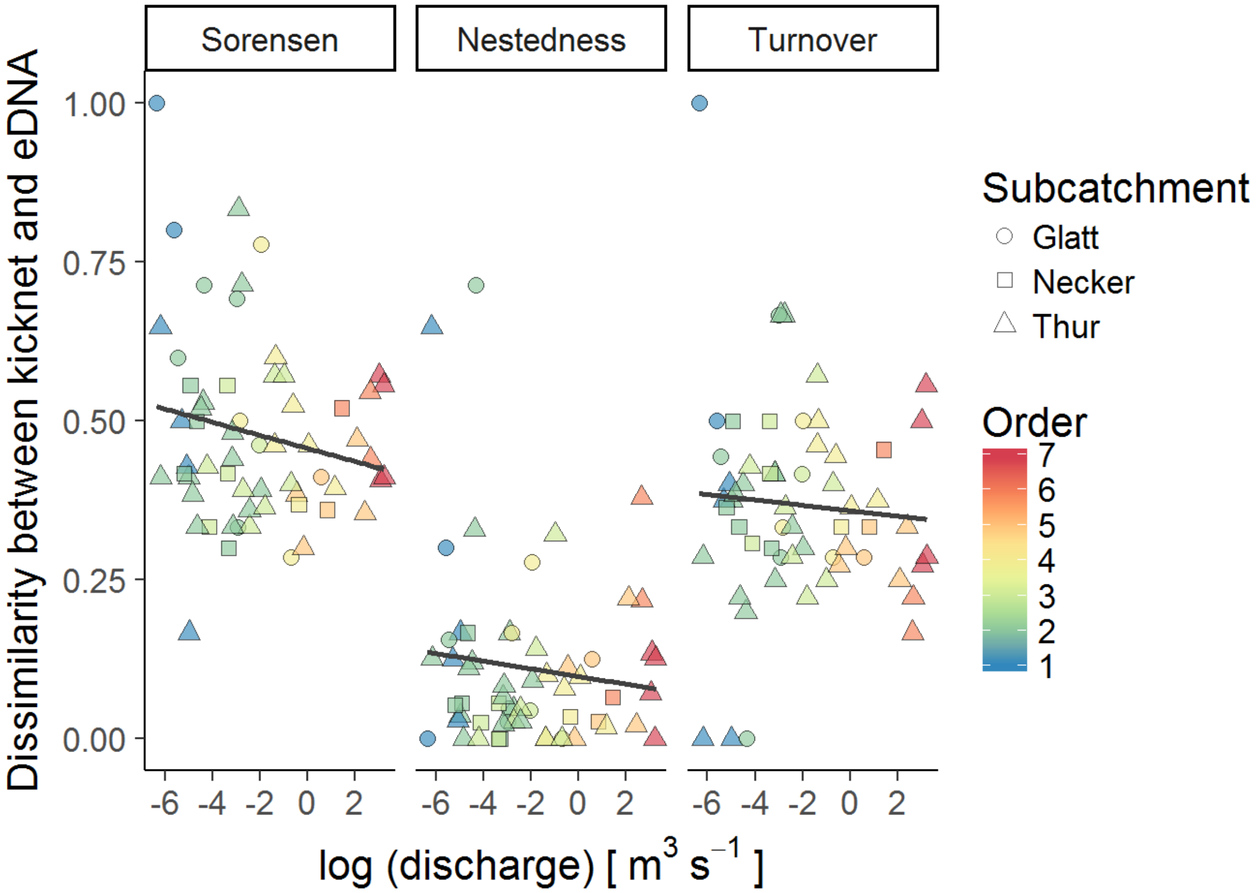


**Fig. S4:** Dissimilarity between kicknet and eDNA samples versus the logarithmic annual mean discharge per site. Inequality is based on Sorensen dissimiliarity and split into nestedness and turnover for EPT genera only. Colors are according to stream order and the shape indicates to what subcatchment the site belongs to. Lines indicate a linear regression. Points are minimally jittered due to overlapping cases.
